# Head Stabilization During Standing in People with Persisting Symptoms after Mild Traumatic Brain Injury

**DOI:** 10.1101/850081

**Authors:** Peter C. Fino, Tiphanie E Raffegeau, Lucy Parrington, Robert J Peterka, Laurie A. King

**Affiliations:** University of Utah, Department of Health, Kinesiology, and Recreation, Salt Lake City, UT; Oregon Health Sciences University, Department of Neurology, Portland, OR; National Center for Rehabilitative Auditory Research, VA Portland Health Care System, Portland, OR

**Keywords:** Posture, Balance, Sway, Sensory Integration, Concussion, Stability

## Abstract

Increased postural sway is often observed in people with mild traumatic brain injury (mTBI), but our understanding of how individuals with mTBI control their head during stance is limited. The purpose of this study was to determine if people with mTBI exhibit increased sway at the head compared with healthy controls. People with persisting symptoms after mTBI (*n* = 59, 41 women) and control participants (*n* = 63, 38 women) stood quietly for one minute in four conditions: eyes open on a firm surface (EO-firm), eyes closed on a firm surface (EC-firm), eyes open on a foam pad (EO-foam), and eyes closed on foam (EC-foam). Inertial sensors at the head, sternum, and lumbar region collected tri-axial accelerations. Root-mean-square (RMS) accelerations in anteroposterior (AP) and mediolateral (ML) directions. Sway ratios between the head and sternum, head and lumbar, and sternum and lumbar region, were compared between groups. Temporal coupling of anti-phase motion between the upper and lower body angular accelerations was assessed with magnitude squared coherence and cross-spectral phase angles. People with mTBI demonstrated greater sway than controls across conditions and directions. During foam-surface conditions, the control group, but not the mTBI group, reduced ML sway at their head and trunk relative to their lumbar by increasing the expression of an anti-phase hip strategy within the frontal plane. These results are consistent with suggestions of inflexible or inappropriate postural control in people with mTBI.

## INTRODUCTION

Considerable work has demonstrated the importance of head stabilization in space (i.e., maintaining static head position within a global reference system) for whole-body postural control during stance (Lund and Broberg, 1983;Di Fabio and Emasithi, 1997;Paloski et al., 2006;Hansson et al., 2010) and locomotion (Grossman et al., 1988;Pozzo et al., 1990;Pozzo et al., 1991;Farkhatdinov et al., 2019). Objective assessments of postural sway in people with mild traumatic brain injury (mTBI) often reveal increased sway of the center of mass (CoM) or center of pressure (CoP) compared with healthy controls (Guskiewicz, 2011;Haran et al., 2016;King et al., 2017;Gera et al., 2018), but these results do not inform us about how individuals may stabilize the head in space. While there is some evidence that head stabilization may be impaired while walking in those with mTBI (Sessoms et al., 2015), we are unaware of studies that specifically examine the sway of the head in individuals with mTBI during standing.

The lack of knowledge about head stability during standing in those with mTBI likely stems from two sources: 1) the predominant use of force plates or sensors at the waist to quantify postural sway; and 2) the reliance on single-link inverted pendulum mechanics to understand postural control. Force plates are ideal for quantifying the center-of-pressure (CoP) and reflective of CoM sway measured with inertial sensors at the waist (Mancini et al., 2012). However, the motion of the CoP and CoM does not match the motion of the head (Sakaguchi et al., 1995) because upright postural control is not strictly determined by single-link inverted pendulum mechanics. While a single-link inverted pendulum representation can account for some aspects of body motion during stance in controlled and unperturbed conditions (Gage et al., 2004), a multi-link representation of body mechanics provides a larger feasible control manifold (Kuo and Zajac, 1993;Hsu et al., 2007). Multi-link postural control can also facilitate adaptations to situational demands (Horak and Nashner, 1986) and sensory integration (Mergner and Rosemeier, 1998;Stoffregen et al., 2000) through in-phase and anti-phase motion between the upper and lower body (Creath et al., 2005). Torque generated at the hip joint can stabilize superior body segments such as the head to optimize the sensitivity of sensory inputs to vestibular and visual signals (Pozzo et al., 1995;Farkhatdinov et al., 2019). In addition to the ankle and hip joint, multi-link postural control models can include a joint at the neck (Nicholas et al., 1998). Neck afferent signals are involved in perceptual functions of self-motion (Pettorossi and Schieppati, 2014) and reflex responses such as the cervico-spinal and cervico-ocular reflex, suggesting somatosensory cervical input converges with vestibular input to mediate multisensory control of orientation, gaze, and posture (Brandt and Bronstein, 2001). Head stabilization could therefore be achieved by anti-phase strategies at the hip, neck, or both.

While numerous studies have investigated postural stability during standing in people with mTBI (Guskiewicz, 2011;Sosnoff et al., 2011;Buckley et al., 2016;Fino et al., 2016;Gera et al., 2018), the control of head stability after mTBI remains unclear. Thus, we conducted an ancillary analysis of a larger study (Fino et al., 2017) to address two pertinent questions: 1) Do individuals with persisting symptoms after mTBI exhibit greater sway at the head compared with healthy controls when standing under various sensory conditions?, and 2) Does the relationship between sway at the lumbar spine level (approximate CoM), sternum, and head differ between those with persisting symptoms after mTBI and healthy controls as sensory conditions change? In accordance with previous reports of larger CoP and CoM sway after mTBI, we hypothesized that individuals with mTBI would sway more at the head compared with controls. We did not have a direction-specific *a priori* hypothesis for the second exploratory question, but we anticipated that segmental sway patterns in people with mTBI would differ from controls under more challenging sensory conditions. For example, if normalized to anthropometry, smaller sway at the head relative to the sternum would indicate more active anti-phase neck control, and smaller sway at the sternum relative to the lumbar would indicate more active anti-phase hip control.

## MATERIALS and METHODS

### Participants

Fifty-nine individuals with chronic (>3 months) balance complaints after a clinically diagnosed mTBI and 63 healthy controls (Table 1) were recruited as part of a larger study (ClinicalTrials.gov Identifier NCT02748109) in people with chronic mTBI (detailed elsewhere; see (Fino et al., 2017)). Briefly, participants were recruited through posters in athletic facilities, physical therapy clinics, hospitals, concussion clinics, community notice boards and cafes in and around the Portland, Oregon metropolitan area. Participants were included if they were: >3 months post-mTBI and reporting a nonzero symptom score on the Sport Concussion Assessment Tool 3 (SCAT3)(Guskiewicz et al., 2013) balance problems symptom, or had no history of brain injury in the past year for the control group; zero to minimal cognitive deficits as determined by the Short Blessed Test (Katzman et al., 1983) (score ≤8); and were between the ages of 18 and 60 years. For the purposes of this study, mTBI was defined and classified using the criteria from the United States Department of Defense: no CT scan, or a normal CT scan if obtained; loss of consciousness not exceeding 30 min; alteration of consciousness/mental state up to 24 h; and post-traumatic amnesia not exceeding one day (Woodson, 2015). The mechanism of injury was not restricted. Exclusion criteria for both groups consisted of: musculoskeletal injury in the previous year that could have seriously impacted gait or balance; current moderate or severe substance abuse; any past peripheral vestibular or oculomotor pathology from before their reported mTBI; or refusal to abstain from medications that could impact their mobility for the duration of testing. Participants were asked to abstain from medications that could impact their mobility starting 24 hours prior to their first testing date. These medications included sedatives, benzodiazepines, narcotic pain medications and alcohol. All recruitment procedures were approved by the Oregon Health & Science University and Veterans Affairs Portland Health Care System joint institutional review board and participants provided written informed consent prior to commencing the study.

**Table 1.** Demographic information for each group of participants. Data are presented as mean (SD) unless otherwise noted.

|  | **mTBI** | **Control** |
| --- | --- | --- |
| **N** | 59 | 63 |
| **Age (yrs)** | 39.3 (10.9) | 37.7 (12.8) |
| **Gender (F / M)** | 41/18 | 38/25 |
| **Height (cm)** | 170.8 (9.4) | 171.5 (9.5) |
| **Mass (kg)** | 79.8 (19.7) | 75.2 (19.0) |
| **NSI Symptom Score\*** | 37.0 (23.0) | 3.0 (5.0) |
| **One or more lifetime mTBI (n)** | 59 | 5 |
| **Time Since Most Recent mTBI (yrs)\*** | 1.12 (1.95) | 13.28 (11.10) |

\* Shown as median (interquartile range) NSI: Neurobehavioral Symptom Inventory

### Procedures

Subjects completed demographic and symptom-related questionnaires (Neurobehavioral Symptom Inventory, NSI(Cicerone and Kalmar, 1995)) and an assessment of postural control for 60 seconds standing under four different sensory conditions: (1) firm ground with eyes open (EO-Firm), (2) firm ground with eyes closed (EC-Firm), (3) foam surface (Airex Balance-pad Elite, Airex AG, Sins, Switzerland) with eyes open (EO-Foam), (4) and foam surface with eyes closed (EC-Foam)(Shumway-Cook and Horak, 1986;Fino et al., 2017;Freeman et al., 2018;Gera et al., 2018). For all conditions, participants were instructed to stand with their feet together with their hands on their hips, and to hold that position for 60 seconds. The duration of 60 seconds was chosen to ensure a long enough period to capture reliable sway data after removing transient effects (Scoppa et al., 2013). If the participant lost balance or deviated from the starting position (e.g., opening eyes during an eyes closed condition) before the completion of the 60 seconds, the trial was stopped. Trials that were stopped early were not repeated, and these trials were excluded from future analysis. Trials were always presented in the same order: EO-Firm, EC-Firm, EO-Foam, EC-Foam, and rest breaks were provided between each condition as needed.

### Data Analysis

Data were collected at 128 Hz using wearable inertial sensors (Opal, APDM Inc.) affixed over the lumbar spine region (approximate CoM), sternum, forehead, and bilaterally on the dorsum of the feet using elastic straps. Each sensor provided tri-axial acceleration, angular velocity, and magnetometer data. For this ancillary analysis, only data from the head, sternum, and lumbar sensors were analyzed.

For each condition, the sensors’ axes were rotated to align with the global coordinate system (Moe-Nilssen, 1998). Acceleration data were low-pass filtered using a phaseless 4^th^ order, 10 Hz low pass Butterworth filter. After filtering, the first and last 10 seconds of each trial were removed to reduce the influence of transient effects from the start of the trial, movement coincident with the end of the trial, or end distortion from filtering. The middle 40 seconds of each trial were retained for analysis. For each condition, accelerations in the transverse plane were extracted for each sensor (Figure 1), and the root mean square (RMS) of AP and ML accelerations were calculated for the head, sternum, and lumbar sensors. To examine the sway at the head relative to the sway at the lumbar, the ratio of head sway over lumbar sway was calculated in each direction for all trials. To account for the effect of pendulum length on linear accelerations, acceleration sway ratios were normalized based on the height of the sensor. For example, the head-to-lumbar sway ratio was multiplied by the ratio of the height of the lumbar sensor over the height of the head sensor,

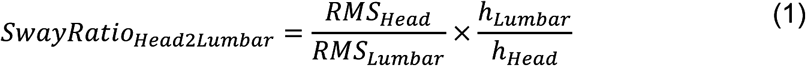

where *h*_*Head*_ and *h*_*Lumbar*_ *are* the height of the head and lumbar sensors, respectively and RMS is the RMS of acceleration at each segment. Similarly, the ratio of head-to-sternum and sternum-to-lumbar sway was also calculated to determine if the neck played a significant role in attenuating acceleration between the lumbar and head. As the height of each sensor was not initially recorded, but the anatomical placement was consistent across subjects, the height of each sensor was estimated based on the percentage of total height based on standard anthropometric data: *h*_*Head*_ = 0.96, *h*_*Sternum*_ = 0.76, *h*_*Lumbar*_ = 0.59 (de Leva, 1996). Sway ratios equal to one indicate single-link sway about the ankle, sway ratios less than 1 indicate anti-phase multi-link sway where the superior segment is stabilized relative to the inferior segment, and sway ratios greater than one indicate in-phase multi-link sway where the superior segment sway is amplified relative to the inferior segment.

**Figure 1.**
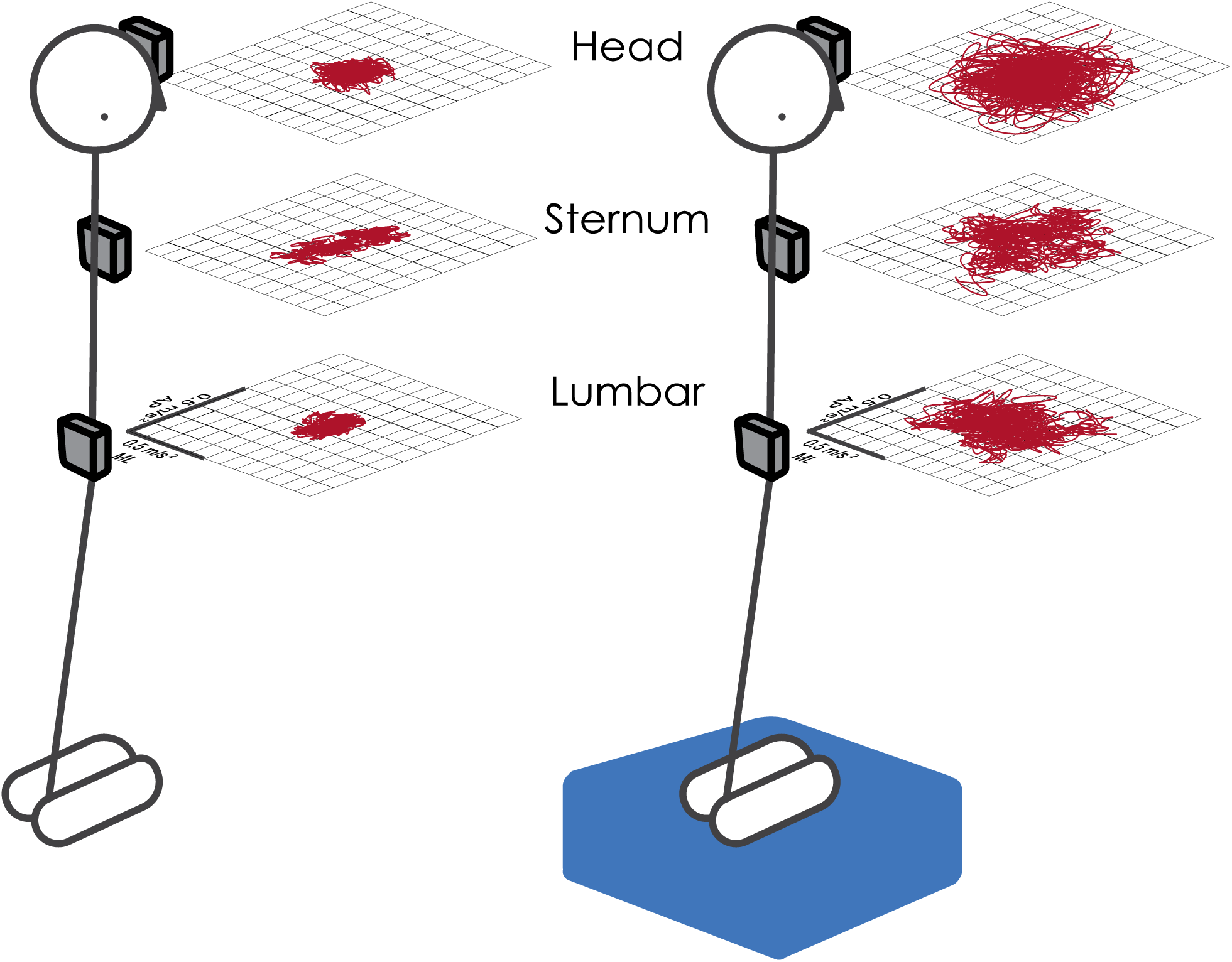
Representation of acceleration sway within the transverse plane at the head, sternum, and lumbar sensor locations for a single subject during EO-Firm (left) and EC-Foam (right) conditions.

To examine the temporal coupling between segmental sway, measurements of AP and ML accelerations at head and lumbar levels were used to calculate angular accelerations of the upper-body trunk/head segment and the lower-body segment within the sagittal and frontal planes, respectively. Cross-spectral analysis between these two segmental angular motions was used to investigate the presence of anti-phase motion coupling between the segments (i.e. 180° out-of-phase motion) and the strength (coherence) of the coupling. Comparisons of segmental angular accelerations to assess temporal coupling between segments is advantageous over the direct use of linear acceleration measures; a perfectly stabilized head would have zero linear acceleration thus preventing any meaningful conclusion about the temporal relation between lumbar and head accelerations, while comparison of segmental angular accelerations would reveal anti-phase temporal coupling.

The angular acceleration at head and lumbar levels with respect to a rotation axis at ground level was calculated by dividing the measured transverse accelerations at these two levels by the respective heights above the ground: α_*head*_ = *a*_*head*_/*h*_*Head*_, α_*L*_ = *a*_*lumbar*_/*h*_*lumbar*_. The upper body angular acceleration α_*UB*_ is given by the difference between α_*head*_ and α_*lumbar*_: α_*UB*_ = α_*head*_ − αα_*lumbar*_ and the lower body angular acceleration α_*LB*_ = α_*lambar*_.

To evaluate the temporal coupling between segments, the magnitude-squared coherence and phase of the cross power spectral density (CSD) of *α*_*UB*_ and *α*_*LB*_ was computed for both AP and ML directions. The coherence and CSD were implemented in MATLAB using the functions ‘mscohere’ and ‘cpsd’, respectively, using a 10 second Hamming window with 50% overlap.

### Statistical Analysis

To examine whether individuals with mTBI and healthy controls had similar rates of failure (e.g., loss of balance before 60 seconds), Chi-squared proportions tests compared the number of failed trials in each condition using a 0.05 significance level. To assess differences between groups across conditions, linear mixed effects regression models were fit for RMS sway at each body segment in each direction. Outcomes were first assessed for normality; all segments and directions exhibited skewed distributions for RMS sway and were therefore log-transformed. Each linear mixed effects regression model contained fixed effects of group, condition, the group×condition interaction, and random intercepts to account for the within-subject correlations. Condition was modeled as a categorical variable with EO-Firm serving as the reference condition. The control group served as the reference condition for group. Post-hoc pairwise comparisons were performed using independent sample t-tests and Cohen’s *d* effect sizes to further investigate group×condition interactions for any sway ratio outcome. As coherence and cross-spectral phase were obtained as complementary secondary outcomes, descriptive statistics were extracted and presented, but no statistical comparisons were performed on these outcomes. All significance values were corrected for multiple comparisons using a false discovery rate (FDR) correction (Benjamini and Hochberg, 1995) and a significance level of 0.05. All statistical analysis was performed in MATLAB (r2018a, The MathWorks Inc.).

## RESULTS

### Task Completion

No participants in the control group failed to complete the 60 seconds during any of the trials. Comparatively, mTBI subjects failed to complete a total of 23 trials (EO-firm: *n* = 2 (3%), *p* = 0.447, EC-firm: *n* = 8 (14%), *p* = 0.008, EO-foam: *n* = 3 (5%), *p* = 0.220, EC-foam: *n* = 10 (17%), *p* = 0.002). All subjects who failed the EO-firm trial failed all other trials. Subjects who failed the EC-firm trial or the EO-foam trial also failed the EC-foam trial. Compared with controls, more mTBI subjects failed the EC trials, but did not have significantly higher failure rates during EO trials.

### Head Sway

Participants with mTBI swayed more at the head for all conditions and directions relative to controls (Tables 2,3). Relative to the baseline EO-Firm condition, RMS head sway increased in both AP and ML directions for every condition in control participants (main condition effects, Table 3). Compared with the between-condition changes in the control group, the mTBI group exhibited larger increases in head sway in both directions and all conditions except for EO-Foam in the ML direction (group×condition interactions, Table 3).

**Table 2.** Descriptive statistics of median and inter-quartile range (IQR) for RMS sway at the head, trunk, and lumbar spine in mTBI and control groups.

| Location | Head |  |  |  | Trunk |  |  |  | Lumbar |  |  |  |
| --- | --- | --- | --- | --- | --- | --- | --- | --- | --- | --- | --- | --- |
| Group | mTBI |  | Control |  | mTBI |  | Control |  | mTBI |  | Control |  |
| Statistic | *Median* | *IQR* | *Median* | *IQR* | *Median* | *IQR* | *Median* | *IQR* | *Median* | *IQR* | *Median* | *IQR* |
| **AP direction** |  |  |  |  |  |  |  |  |  |  |  |  |
| EO-Firm | 0.18 | 0.12 | 0.12 | 0.08 | 0.13 | 0.06 | 0.11 | 0.06 | 0.11 | 0.05 | 0.08 | 0.05 |
| EC-Firm | 0.24 | 0.22 | 0.14 | 0.08 | 0.15 | 0.12 | 0.11 | 0.04 | 0.13 | 0.10 | 0.09 | 0.05 |
| EO-Foam | 0.24 | 0.17 | 0.15 | 0.07 | 0.16 | 0.10 | 0.13 | 0.06 | 0.14 | 0.08 | 0.11 | 0.03 |
| EC-Foam | 0.40 | 0.23 | 0.22 | 0.12 | 0.27 | 0.23 | 0.17 | 0.07 | 0.22 | 0.16 | 0.13 | 0.06 |
| **ML direction** |  |  |  |  |  |  |  |  |  |  |  |  |
| EO-Firm | 0.12 | 0.08 | 0.08 | 0.03 | 0.08 | 0.06 | 0.05 | 0.02 | 0.06 | 0.04 | 0.05 | 0.02 |
| EC-Firm | 0.15 | 0.13 | 0.08 | 0.03 | 0.09 | 0.10 | 0.06 | 0.02 | 0.08 | 0.06 | 0.05 | 0.02 |
| EO-Foam | 0.18 | 0.19 | 0.12 | 0.05 | 0.14 | 0.12 | 0.09 | 0.04 | 0.12 | 0.08 | 0.10 | 0.03 |
| EC-Foam | 0.32 | 0.22 | 0.17 | 0.05 | 0.25 | 0.21 | 0.14 | 0.05 | 0.20 | 0.12 | 0.15 | 0.06 |

*Note: All values given in units of m/s<sup>2</sup>*

**Table 3.** Mixed effect regression model fit and parameters for head, sternum, and lumbar accelerations.

|  | *Head* |  |  | *Sternum* |  |  | *Lumbar* |  |  |
| --- | --- | --- | --- | --- | --- | --- | --- | --- | --- |
| Fixed-Effects | $\beta$ | SE | Corrected *p* | $\beta$ | SE | Corrected *p* | $\beta$ | SE | Corrected *p* |
| **AP direction** |  |  |  |  |  |  |  |  |  |
| Intercept | -2.07 | 0.06 | < **0.001** | -2.26 | 0.06 | < **0.001** | -2.49 | 0.05 | < **0.001** |
| Condition: EO-Firm | **Ref.** | -- | -- | **Ref.** | -- | -- | **Ref.** | -- | -- |
| Condition: EC-Firm | 0.15 | 0.06 | **0.019** | 0.09 | 0.06 | 0.191 | 0.04 | 0.05 | 0.463 |
| Condition: EO-Foam | 0.17 | 0.06 | **0.006** | 0.19 | 0.06 | **0.003** | 0.26 | 0.05 | < **0.001** |
| Condition: EC-Foam | 0.59 | 0.06 | < **0.001** | 0.53 | 0.06 | < **0.001** | 0.54 | 0.05 | < **0.001** |
| Group: Control | **Ref.** | -- | -- | **Ref.** | -- | -- | **Ref.** | -- | -- |
| Group: mTBI | 0.29 | 0.09 | **0.002** | 0.19 | 0.09 | 0.051 | 0.25 | 0.07 | **0.002** |
| Condition: EO-Firm × Group | **Ref.** | -- | -- | **Ref.** | -- | -- | **Ref.** | -- | -- |
| Condition: EC-Firm × Group | 0.25 | 0.08 | **0.008** | 0.19 | 0.08 | **0.041** | 0.25 | 0.08 | **0.004** |
| Condition: EO-Foam × Group | 0.21 | 0.08 | **0.019** | 0.18 | 0.08 | **0.049** | 0.11 | 0.08 | 0.244 |
| Condition: EC-Foam × Group | 0.27 | 0.08 | **0.004** | 0.30 | 0.08 | **0.001** | 0.21 | 0.08 | **0.018** |
| **ML direction** |  |  |  |  |  |  |  |  |  |
| Intercept | -2.55 | 0.05 | < **0.001** | -2.99 | 0.06 | < **0.001** | -3.04 | 0.05 | < **0.001** |
| Condition: EO-Firm | **Ref.** | -- | -- | **Ref.** | -- | -- | **Ref.** | -- | -- |
| Condition: EC-Firm | 0.08 | 0.04 | **0.048** | 0.18 | 0.05 | **0.003** | 0.14 | 0.04 | **0.003** |
| Condition: EO-Foam | 0.44 | 0.04 | < **0.001** | 0.54 | 0.05 | < **0.001** | 0.67 | 0.04 | < **0.001** |
| Condition: EC-Foam | 0.81 | 0.04 | < **0.001** | 1.02 | 0.05 | < **0.001** | 1.15 | 0.04 | < **0.001** |
| Group: Control | **Ref.** | -- | -- | **Ref.** | -- | -- | **Ref.** | -- | -- |
| Group: mTBI | 0.43 | 0.08 | < **0.001** | 0.48 | 0.09 | < **0.001** | 0.38 | 0.07 | < **0.001** |
| Condition: EO-Firm × Group | **Ref.** | -- | -- | **Ref.** | -- | -- | **Ref.** | -- | -- |
| Condition: EC-Firm × Group | 0.17 | 0.06 | **0.016** | 0.13 | 0.08 | 0.189 | 0.08 | 0.06 | 0.302 |
| Condition: EO-Foam × Group | 0.09 | 0.06 | 0.259 | 0.12 | 0.08 | 0.199 | -0.09 | 0.06 | 0.220 |
| Condition: EC-Foam × Group | 0.16 | 0.07 | **0.025** | 0.19 | 0.08 | **0.033** | -0.03 | 0.07 | 0.722 |

*Note:* Degrees of freedom = 456 for all effects. Significance denoted by bolded *p*-value.
Parameter estimates: slope estimate ( $\beta$ ), standard error (SE), and corrected *p*-value (*p*).

### Sternum Sway

Participants with mTBI exhibited greater RMS sway at the sternum for all conditions in the ML direction only (Tables 2,3). Relative to the baseline EO-Firm condition, RMS sternum sway increased in both AP and ML directions for all other conditions in control participants except for EC-Firm in the AP direction (condition effects, Table 3). Compared with the between-condition changes in the control group, the mTBI group had larger increases in AP sternum sway in all conditions and larger changes in ML sternum sway in the EC-Foam condition (group×condition interactions, Table 3).

### Lumbar Sway

Similar to sway at the head, participants with mTBI exhibited greater RMS sway at the lumbar for all conditions and directions (Tables 2,3). Relative to the baseline EO-Firm condition, RMS lumbar sway increased in both directions for all other conditions in control participants except for EC-Firm in the AP direction (condition effects, Table 3). Compared to the between-condition changes relative to the baseline EO-Firm condition in the control group, the mTBI group had larger increases in AP lumbar sway in EC-Firm and EC-Foam conditions (group×condition interactions, Table 3).

### Head to Lumbar Sway Ratio

In foam conditions, control participants decreased the ML head-to-lumbar sway ratio relative to the baseline EO-Firm condition (condition effects, Table 4, Figure 2F). Comparatively, the mTBI group exhibited little change in the ML head-to-lumbar sway ratio in foam conditions (EO-Foam, EC-Foam) relative to the EO-Firm condition (group×condition interactions, Table 4, Figure 2F). Post-hoc t-tests revealed the mTBI group had significantly greater sway ratios than the control group during foam conditions (EO-Foam *d* = 0.74, *adjusted p* < 0.001; EC-Foam *d* = 0.91, *adjusted p* < 0.001). No effects of group, condition, or interactions were detected for the head-to-lumbar sway ratio in the AP direction.

**Table 4.**
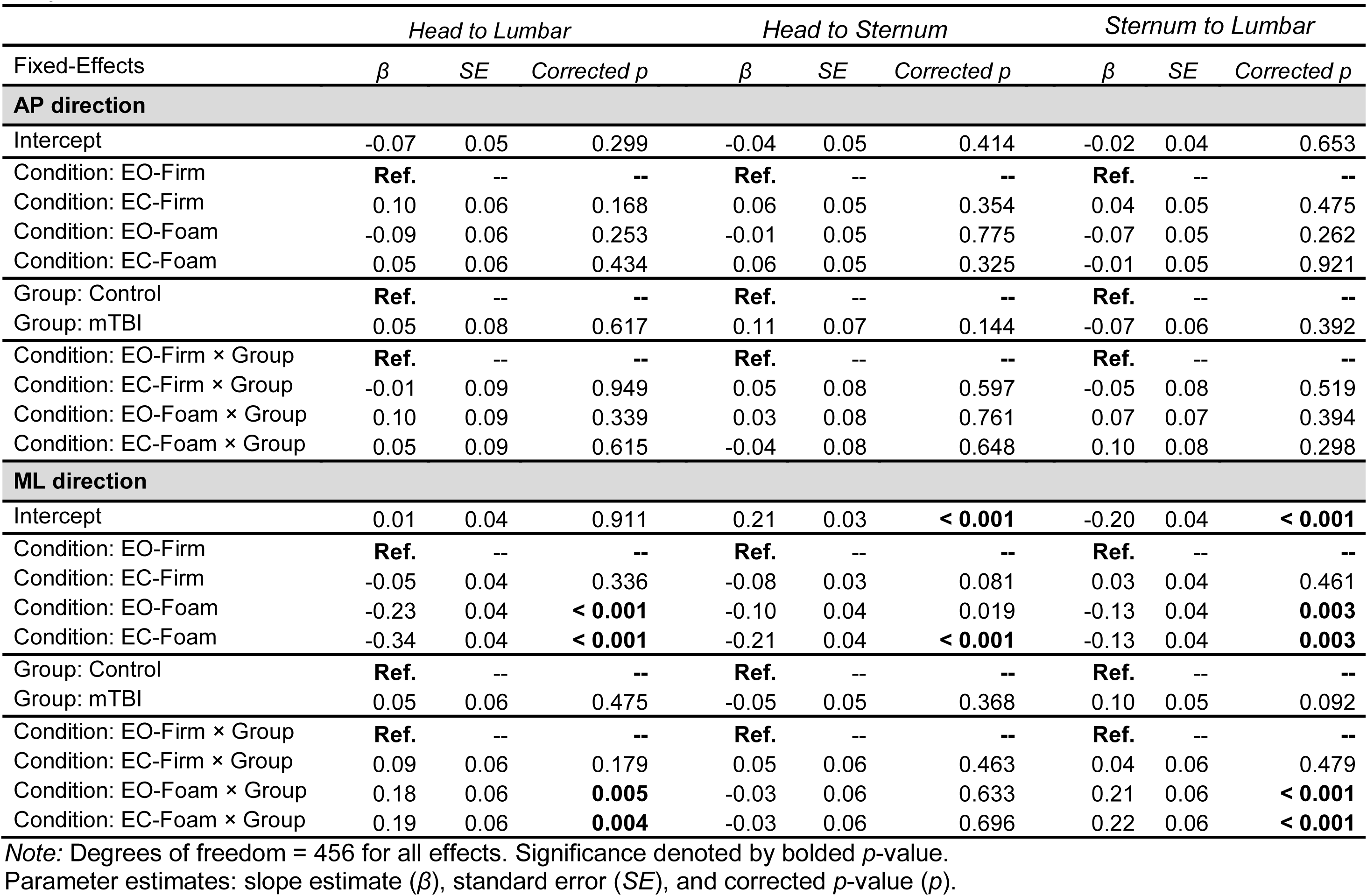
Mixed effect regression model fit and parameters for head to lumbar, head to sternum, and sternum to lumbar sway ratios.

### Head to Sternum Sway Ratio

The head-to-sternum sway ratio did not differ by group or condition in the AP direction. In the ML direction, the head-to-sternum sway ratio decreased in the EC-Foam condition only (condition effects, Table 4); no other group, condition, or group×condition effects were found.

### Sternum to Lumbar Sway Ratio

In foam conditions, control participants decreased the ML sternum-to-lumbar sway ratio relative to the baseline EO-Firm condition (condition effects, Table 4). Comparatively, the mTBI group exhibited increased ML sternum-to-lumbar sway ratio in foam conditions (EO-Foam, EC-Foam) (group×condition interactions, Table 4). Post-hoc t-tests revealed the mTBI group had significantly greater sway ratios than the control group during foam conditions (EO-Foam *d* = 1.05, *adjusted p* < 0.001; EC-Foam *d* = 1.15, *adjusted p* < 0.001). No effects of group, condition, or interactions were found for the sternum-to-lumbar sway ratio in the AP direction.

**Figure 2.**
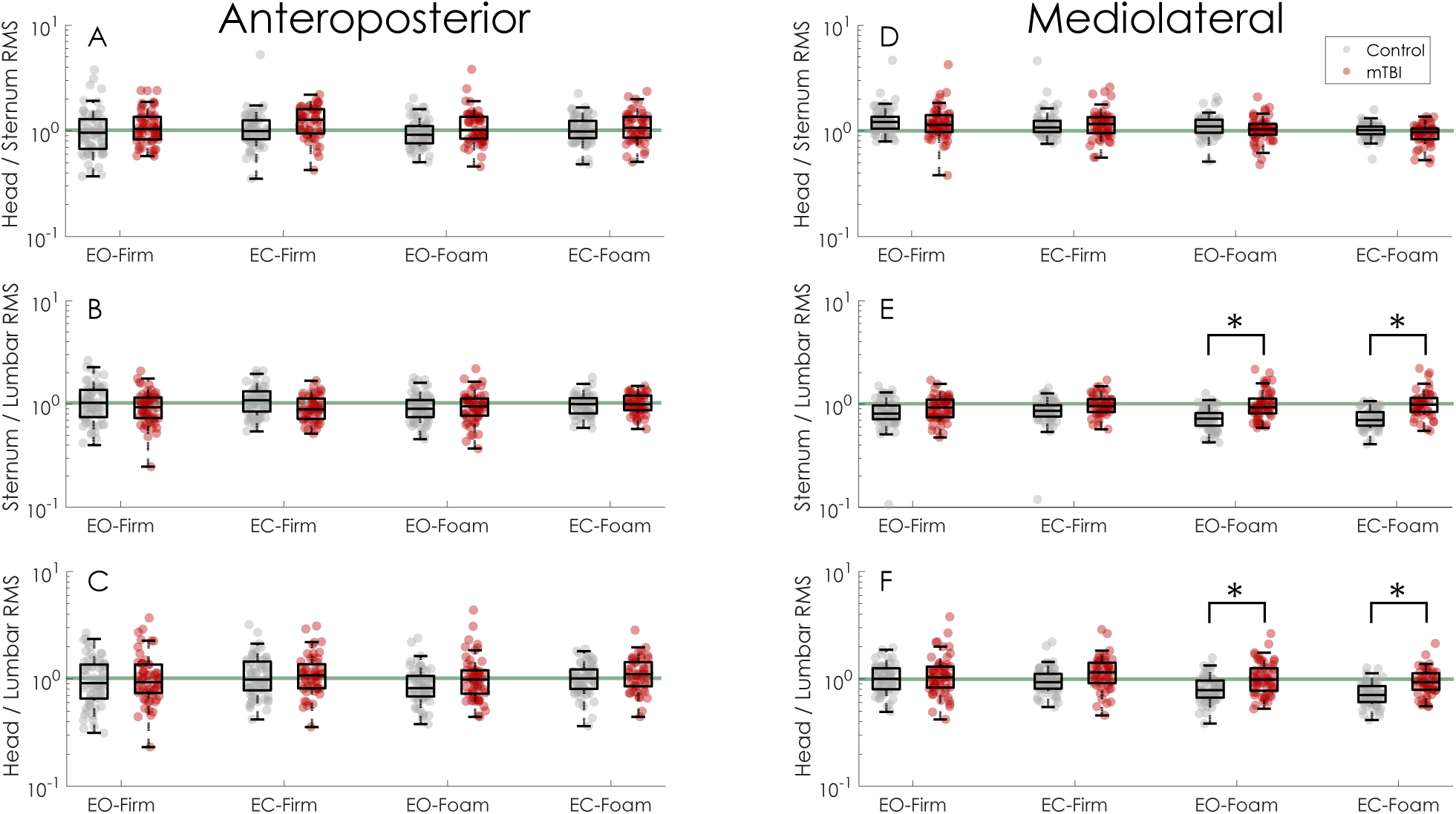
Sway ratios for each direction and condition. A-C) Sway ratios for the head-to-sternum, sternum-to-lumbar, and head-to-lumbar sway ratios in the anteroposterior direction. D-F) Sway ratios for the head-to-sternum, sternum-to-lumbar, and head-to-lumbar sway ratios in the mediolateral direction. Across all figures, the horizontal green line indicates a sway ratio equal to one. In E and F, * indicates significant between-group differences based on post-hoc pairwise t-tests.

### Temporal Coupling of Segments

The coherence and cross-spectral phase between *α*_*UB*_ and *α*_*LB*_ in the frequency band most associated with postural sway (<1 Hz) were consistent with the sway ratio resultss. In foam conditions relative to firm conditions, control participants exhibited stronger anti-phase control of the upper and lower body segments in the frontal plane, characterized by greater coherence between frontal plane *α*_*UB*_ and *α*_*LB*_ and a cross-spectral phase angle near 180° (Figure 3). Despite having similar coherence and cross-spectral phase to controls in firm-ground conditions, the mTBI group exhibited a weaker anti-phase relationship in foam conditions, characterized by lower coherence and cross-spectral phase angles that were farther from 180°. In both groups, a peak in coherence was observed between 0-1 Hz, centered between 0.35-0.50 Hz, during the foam conditions. Consistent with the sway ratio results, no group differences were observed in coherence or cross-spectral phase within the sagittal plane (Figure 4).

**Figure 3.**
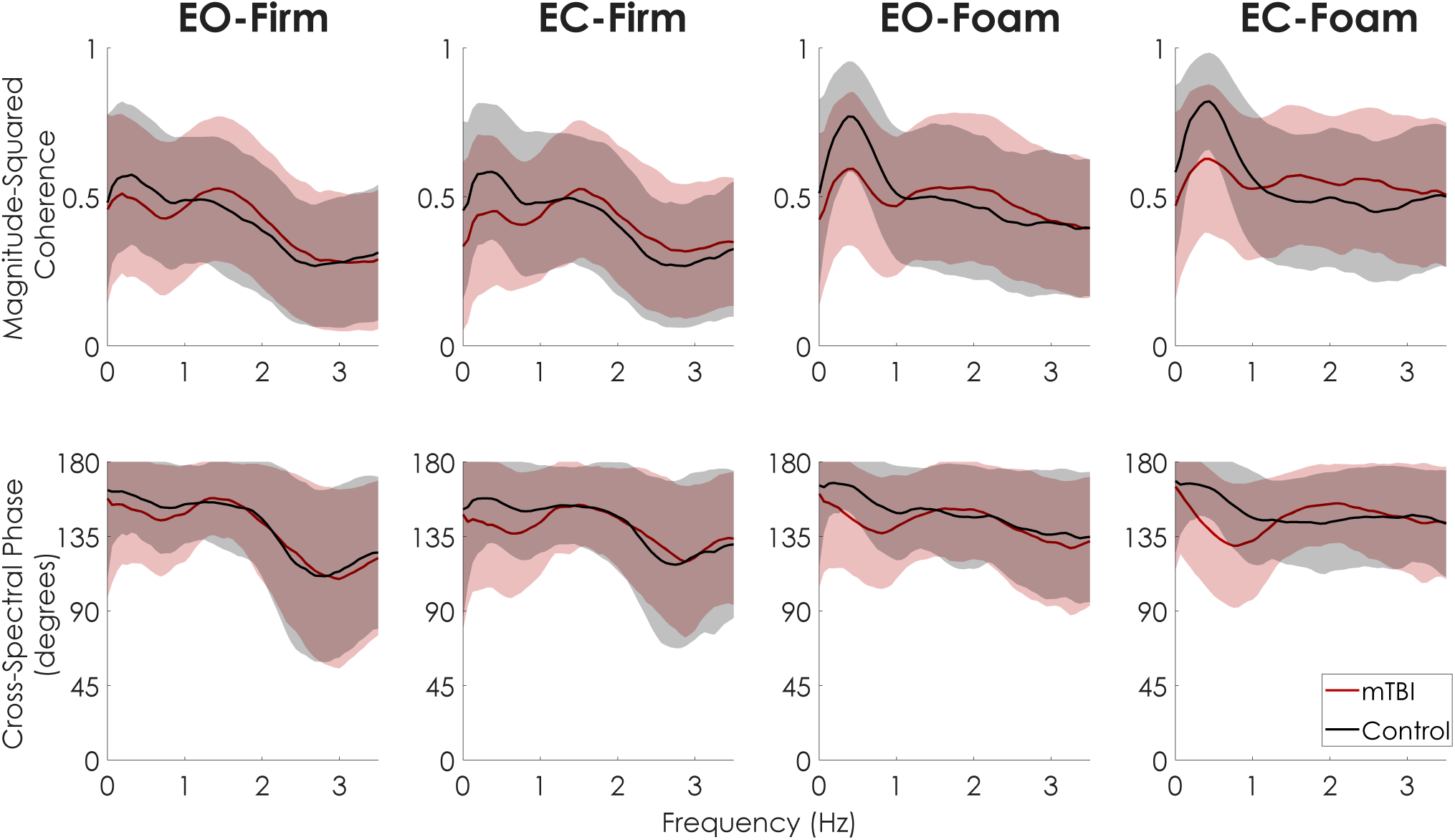
Mean and standard deviation envelopes (±one SD) of magnitude-squared coherence (top) and cross-spectral phase (bottom) between frontal plane *α*_*UB*_ and *α*_*LB*_ for both mTBI (red) and Control (black) groups in each condition. In the foam conditions, the Control group exhibited greater coherence between 0-1 Hz, and maintained a stronger anti-phase relationship between *α*_*UB*_ and *α*_*LB*_ compared to the mTBI group. The stronger anti-phase relationship are consistent with increased head-in-space stabilization in the foam conditions.

**Figure 4.**
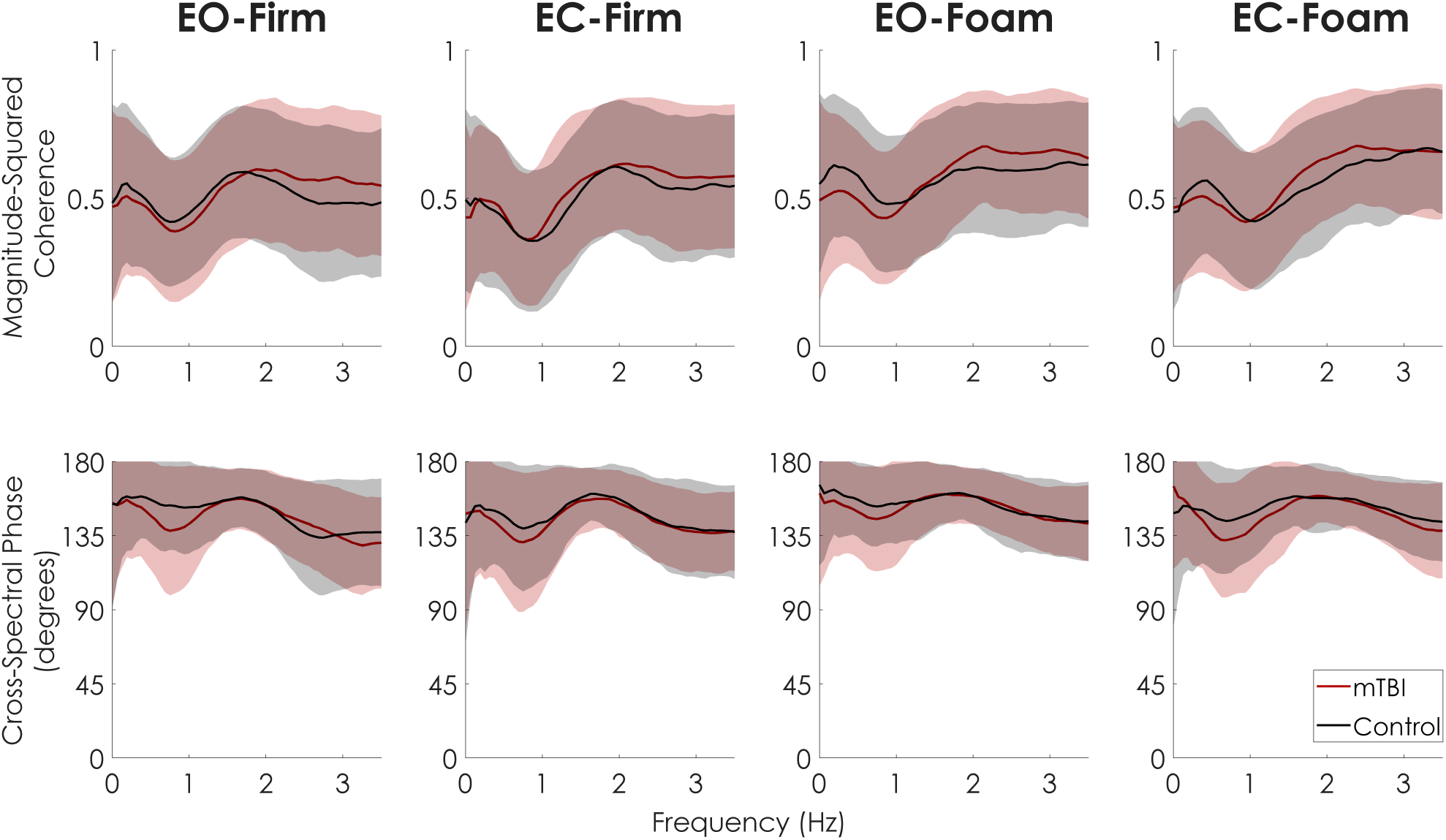
Mean and standard deviation envelopes (±one SD) of magnitude-squared coherence (top) and cross-spectral phase (bottom) between sagittal plane *α*_*UB*_ and *α*_*LB*_ for both mTBI (red) and Control (black) groups in each condition. No group differences are noted.

## DISCUSSION

Our primary objective was to compare postural sway at the head between individuals with mTBI and healthy control subjects under varying sensory conditions. Additionally, we sought to examine the relationship between stabilization of the head and stabilization of inferior segments (sternum and lumbar) to understand potential postural control strategies. We found individuals with mTBI swayed more at the head, sternum, and lumbar spine, extending previous results indicating postural sway of the CoM or CoP is greater in people with mTBI (Guskiewicz, 2011;King et al., 2017;Gera et al., 2018). Between-group differences in RMS sway, however, were greatest at the head location. Additionally, the largest between-group differences were observed in the ML direction, agreeing with previous studies suggesting the importance of accelerometry-derived ML sway in differentiating acute, symptomatic mTBI from healthy controls (King et al., 2017).

We found healthy control subjects tended to stabilize the head in space when standing on foam, while individuals with mTBI did not exhibit the same degree of head stabilization. Fujisawa et al. (2005) represented body mechanics as a double inverted pendulum that allowed joint rotations at the ankles and hip while assuming a rigid head-on-trunk coupling and showed that balance control shifted toward greater use of hip joint torques compared to ankle joint torques when the surface was narrowed. With double pendulum body mechanics, proprioceptors at the ankle determine the ankle angle, proprioceptors at the hip determines the hip angle, and the visual and vestibular sensors determine the angle of the head and trunk relative to vertical (Figure 5). At any point in time, the horizontal position of the CoM, *x*_*CoM*_ can then be described using one of three equations

**Figure 5.**
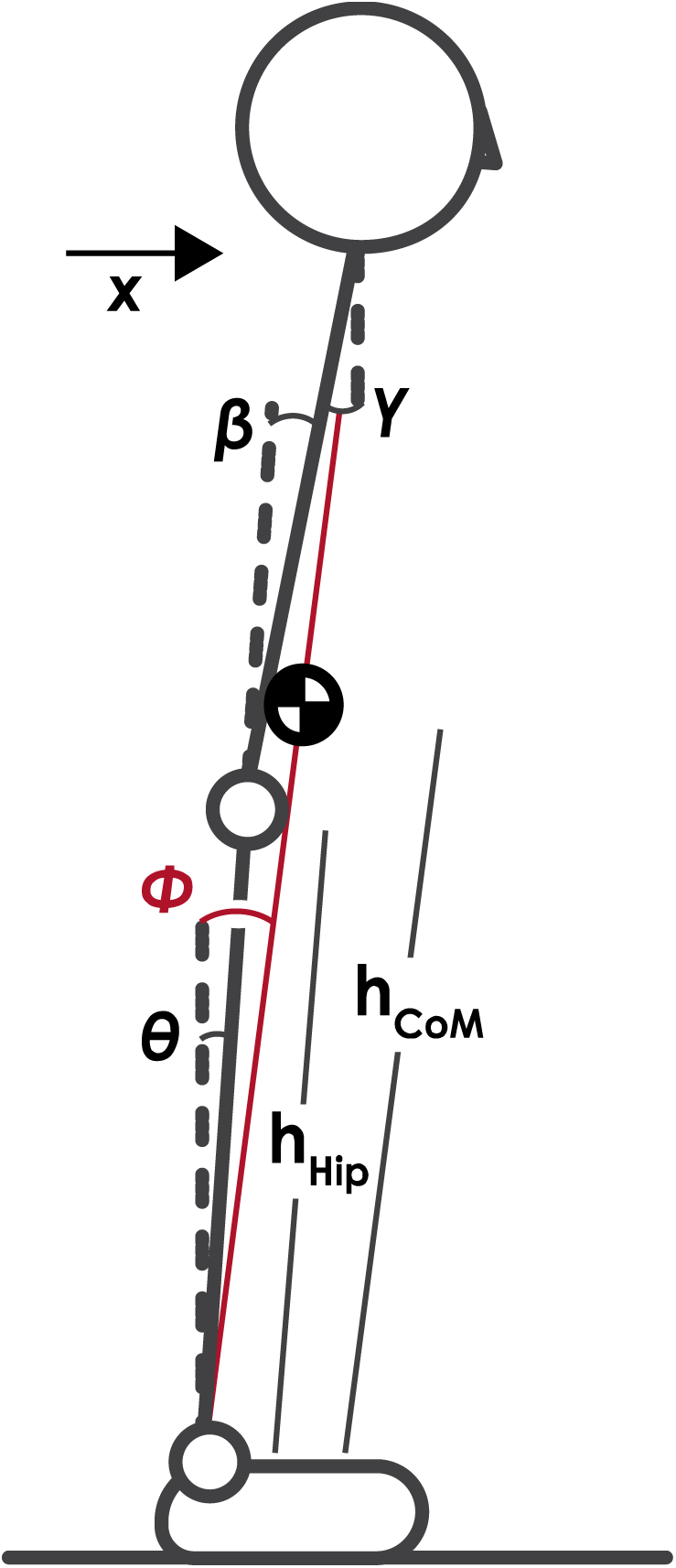
Schematic of two-link postural control state estimation. Motion in the sagittal or frontal plane can be determined from the combined height of the hip joint, *h*_*hip*_, height of the center of mass, *h*_*CoM*_, and two of the three angles: the angle of the ankle with respect to vertical, *θ*, encoded through ankle proprioceptors; the angle of the hip with respect to the lower body, *β*, encoded through hip proprioceptors; and the angle of the upper body with respect to vertical, *γ*, encoded through visual and vestibular sensory systems. Using Equations 6, 7, or 8, the horizontal position of the CoM and the angle of a single-link inverted pendulum model, *Φ*, (shown in red) can be determined. All angles are defined as positive for a counterclockwise rotation with respect to the reference.

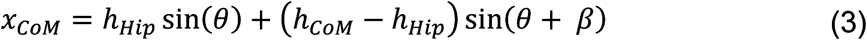

where *h*_*Hip*_ is the height of the hip, *h*_*CoM*_ is the height of the CoM, *θ* is the ankle angle, and *β* is the hip angle,

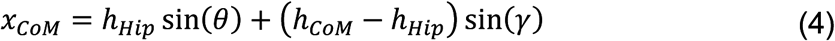

where *γ* is the angle of the upper body relative to vertical, or

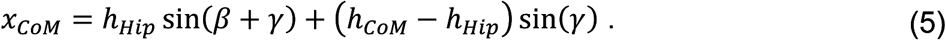

Adding a noise term onto each angle estimate to reflect sensory noise common and dividing by *h*_*CoM*_ yields

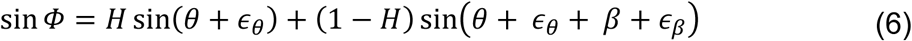

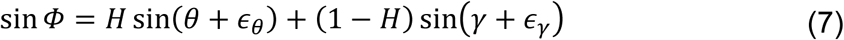

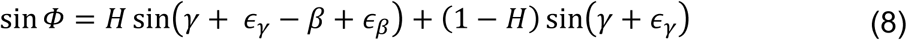

where *Φ* is the angle with respect to vertical of a single inverted pendulum connecting the CoM to the ankle joint and *H* is the ratio 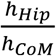. Since *H* is routinely close to 0.9, the above equations are largely dominated by the first term. Therefore, the state estimation of the position of the center of mass will be influenced by the sensory error at the ankle *ϵ*_α_ in Eq.6 and 7, or the combined sensory error at the hip and head *ϵ*_*β*_ *+ ϵ*_*γ*_ in Eq. 8. The head-in-space stabilization demonstrated by controls may have been achieved by a strategy that places greater reliance on hip proprioceptors and visual/vestibular information at the head to estimate body position (Eq. 8) when ankle proprioceptive cues are unreliable in the foam surface condition. The reduced head stabilization in the mTBI group may be due to an inflexible postural control strategy that was unable to shift toward increased use of hip and head sensory information to reduce the use of unreliable ankle proprioceptive cues. This interpretation agrees with studies using entropic measures of sway; individuals with mTBI, at various times since injury, have greater regularity and less complexity in their postural sway that are typically indicative of inflexible postural control (Cavanaugh et al., 2006;Sosnoff et al., 2011;Buckley et al., 2016;Fino et al., 2016;Schmidt et al., 2018). However, the postural control strategy exhibited by the mTBI group may not be inflexible, but rather inappropriate given the task demands; a perception of instability or fear of falling related to self-reported symptoms may increase muscle co-contraction at the hips thereby decreasing head stabilization through anti-phase motion about the hip. Future electromyography studies focused on the muscles that span the hip joint are needed to directly answer this question.

Our results also suggest the head-in-space stabilization exhibited by control subjects during foam conditions was achieved primarily through an anti-phase relationship between the angular acceleration of the upper and lower body due to control at the hip. While ankle and hip strategies are traditionally defined in the sagittal plane (Horak and Nashner, 1986), similar control mechanisms consisting of in-phase (i.e., ankle strategy) and anti-phase (i.e., hip strategy) motion of the upper and lower body exist in the frontal plane (Zhang et al., 2007;Goodworth and Peterka, 2010) - we only detected this head stabilization and increased hip strategy during foam conditions in the ML direction. The ML sternum-to-lumbar sway ratio was less than one for all conditions in healthy controls, with lower sway ratios in foam conditions (see Figure 2E). Additionally, the coherence between upper and lower body frontal-plane angular accelerations increased in the foam conditions while maintaining a cross-spectral phase angle near 180°, indicating a stronger anti-phase relationship in the dominant frequency band of postural sway (Zatsiorsky and Duarte, 1999). These results are consistent with the idea that a continuum exists between ankle and hip strategies (Creath et al., 2005). Specific to the narrow stance width employed here, ML balance control can shift from an in-phase ankle strategy to an anti-phase hip strategy based on the amplitude of surface perturbations (Goodworth and Peterka, 2010). Thus, the increased expression of an anti-phase hip strategy in the ML direction when standing on the foam surface in healthy control subjects may be specific to the narrow stance width used here.

The ML head-to-sternum ratios were greater than or equal to one for all conditions, indicating a lack of a strong anti-phase relationship in both groups (see Figure 2D). These results suggest that the stiffness of the neck joint increased in the EC-Foam trial, but no active stabilization caused by anti-phase motion was observed at the neck. Pozzo et al. noted head stabilization in the frontal plane during complex balance tasks tended to occur at the neck, but a rigid head-trunk unit with minimal actuation at the neck compensated for small oscillations (Pozzo et al., 1995). Our results suggesting both groups increased the head-to-trunk coupling during EC-Foam conditions are consistent with the small oscillation condition reported by Pozzo et al. (1995). It is possible that complex balance tasks with larger frontal plane angular displacements than those examined here may elicit compensation utilizing head-on-trunk stabilization that reveals differences between individuals with mTBI and healthy controls. Nevertheless, the lack of a between-group difference in the head-to-sternum sway ratio suggests that neck problems in mTBI subjects, such as neck pain or whiplash, did not influence our results.

Several limitations should be acknowledged and considered. First, participants were tested with their shoes on. Traditionally, postural sway would be assessed barefoot to remove any confounding factor of footwear. However, out of concern for safety and other aspects of the larger study, balance assessments were performed with shoes. Therefore, some heterogeneity in our data may stem from differences in footwear. Second, the actual height of the sensors was not recorded at the time of data collection. As the anatomical placement of the sensors was consistent across subjects, we relied on anthropometric ratios to estimate the height of the sensors. However, as anthropometry varies across individuals, the use of ratios likely introduced some degree of random error into our results. Nevertheless, our findings were robust and it is unlikely this error was biased based on group. Finally, we excluded trials in which individuals lost balance before the end of the trial, removing 17% of our mTBI subjects from the EC-Foam condition and potentially leading to a bias in our sample. However, excluding incomplete trials was a conservative approach. A loss of balance creates extremely large RMS values. Since these trials only occurred in our mTBI groups, removing these trials with a loss of balance likely led to smaller sway in the mTBI group and may have biased our results toward smaller between-group effects. Our results should therefore be interpreted as the conservative estimate of the difference in head stability between mTBI and controls.

In conclusion, we found that people with persisting balance complaints following mTBI exhibited greater sway, quantified with inertial sensors on the head, sternum, and lumbar, compared with healthy control subjects. Further, control subjects reduced the sway at the head relative to sway at the lumbar when standing on foam by increasing the strength of the anti-phase relationship of upper body and lower body angular acceleration. Conversely, those with mTBI did not change postural control strategies between conditions. The attenuation of sway predominantly occurred over the trunk segment, suggestive of a shift towards increasing expression of an anti-phase hip strategy that was supported by coherence and cross-spectral phase results. Speculatively, these results suggest healthy control subjects are more capable, or more willing, to shift control into head-centric postural control using hip torque, while people with persisting symptoms after mTBI may continue to use the same strategy regardless of sensory information.

## ACKNOWLEDGEMENTS

The authors would like to acknowledge the research assistants who contributed to the subject recruitment, enrollment, and data collection throughout this process. This work was supported by the Assistant Secretary of Defense for Health Affairs under Award No. W81XWH-15-1-0620. Additionally, PCF was supported by the Eunice Kennedy Shiver National Institute of Child Health & Human Development of the National Institutes of Health under Award Number K12HD073945. Opinions, interpretations, conclusions and recommendations are those of the authors and are not necessarily endorsed by the Department of Defense or the official views of the National Institutes of Health.

## References

Benjamini, Y., and Hochberg, Y. (1995). Controlling the False Discovery Rate - a Practical and Powerful Approach to Multiple Testing. Journal of the Royal Statistical Society Series B-Statistical Methodology 57, 289–300.

Brandt, T., and Bronstein, A.M. (2001). Cervical vertigo. J Neurol Neurosurg Psychiatry 71, 8–12.

Buckley, T.A., Oldham, J.R., and Caccese, J.B. (2016). Postural control deficits identify lingering post-concussion neurological deficits. J Sport Health Sci 5, 61–69.

Cavanaugh, J.T., Guskiewicz, K.M., Giuliani, C., Marshall, S., Mercer, V.S., and Stergiou, N. (2006). Recovery of postural control after cerebral concussion: new insights using approximate entropy. J Athl Train 41, 305–313.

Cicerone, K.D., and Kalmar, K. (1995). Persistent postconcussion syndrome: The structure of subjective complaints after mild traumatic brain injury. The Journal of Head Trauma Rehabilitation 10, 1–17.

Creath, R., Kiemel, T., Horak, F., Peterka, R., and Jeka, J. (2005). A unified view of quiet and perturbed stance: simultaneous co-existing excitable modes. Neurosci Lett 377, 75–80.

De Leva, P. (1996). Adjustments to Zatsiorsky-Seluyanov’s segment inertia parameters. J Biomech 29, 1223–1230.

Di Fabio, R.P., and Emasithi, A. (1997). Aging and the mechanisms underlying head and postural control during voluntary motion. Phys Ther 77, 458–475.

Farkhatdinov, I., Michalska, H., Berthoz, A., and Hayward, V. (2019). Gravito-inertial ambiguity resolved through head stabilization. Proc Math Phys Eng Sci 475, 20180010.

Fino, P.C., Nussbaum, M.A., and Brolinson, P.G. (2016). Decreased high-frequency center-of-pressure complexity in recently concussed asymptomatic athletes. Gait Posture 50, 69–74.

Fino, P.C., Peterka, R.J., Hullar, T.E., Murchison, C., Horak, F.B., Chesnutt, J.C., and King, L.A. (2017). Assessment and rehabilitation of central sensory impairments for balance in mTBI using auditory biofeedback: a randomized clinical trial. BMC Neurol 17, 41.

Freeman, L., Gera, G., Horak, F.B., Blackinton, M.T., Besch, M., and King, L. (2018). Instrumented Test of Sensory Integration for Balance: A Validation Study. J Geriatr Phys Ther 41, 77–84.

Fujisawa, N., Masuda, T., Inaoka, Y., Fukuoka, H., Ishida, A., and Minamitani, H. (2005). Human standing posture control system depending on adopted strategies. Med Biol Eng Comput 43, 107–114.

Gage, W.H., Winter, D.A., Frank, J.S., and Adkin, A.L. (2004). Kinematic and kinetic validity of the inverted pendulum model in quiet standing. Gait Posture 19, 124–132.

Gera, G., Chesnutt, J., Mancini, M., Horak, F.B., and King, L.A. (2018). Inertial Sensor-Based Assessment of Central Sensory Integration for Balance After Mild Traumatic Brain Injury. Mil Med 183, 327–332.

Goodworth, A.D., and Peterka, R.J. (2010). Influence of stance width on frontal plane postural dynamics and coordination in human balance control. J Neurophysiol 104, 1103–1118.

Grossman, G.E., Leigh, R.J., Abel, L.A., Lanska, D.J., and Thurston, S.E. (1988). Frequency and velocity of rotational head perturbations during locomotion. Exp Brain Res 70, 470–476.

Guskiewicz, K.M. (2011). Balance assessment in the management of sport-related concussion. Clin Sports Med 30, 89-102, ix.

Guskiewicz, K.M., Register-Mihalik, J., Mccrory, P., Mccrea, M., Johnston, K., Makdissi, M., Dvorak, J., Davis, G., and Meeuwisse, W. (2013). Evidence-based approach to revising the SCAT2: introducing the SCAT3. Br J Sports Med 47, 289–293.

Hansson, E.E., Beckman, A., and Hakansson, A. (2010). Effect of vision, proprioception, and the position of the vestibular organ on postural sway. Acta Otolaryngol 130, 1358–1363.

Haran, F.J., Slaboda, J.C., King, L.A., Wright, W.G., Houlihan, D., and Norris, J.N. (2016). Sensitivity of the Balance Error Scoring System and the Sensory Organization Test in the Combat Environment. J Neurotrauma 33, 705–711.

Horak, F.B., and Nashner, L.M. (1986). Central programming of postural movements: adaptation to altered support-surface configurations. J Neurophysiol 55, 1369–1381.

Hsu, W.L., Scholz, J.P., Schoner, G., Jeka, J.J., and Kiemel, T. (2007). Control and estimation of posture during quiet stance depends on multijoint coordination. J Neurophysiol 97, 3024–3035.

Katzman, R., Brown, T., Fuld, P., Peck, A., Schechter, R., and Schimmel, H. (1983). Validation of a short Orientation-Memory-Concentration Test of cognitive impairment. Am J Psychiatry 140, 734–739.

King, L.A., Mancini, M., Fino, P.C., Chesnutt, J., Swanson, C.W., Markwardt, S., and Chapman, J.C. (2017). Sensor-Based Balance Measures Outperform Modified Balance Error Scoring System in Identifying Acute Concussion. Ann Biomed Eng 45, 2135–2145.

Kuo, A.D., and Zajac, F.E. (1993). Human standing posture: multi-joint movement strategies based on biomechanical constraints. Prog Brain Res 97, 349–358.

Lund, S., and Broberg, C. (1983). Effects of different head positions on postural sway in man induced by a reproducible vestibular error signal. Acta Physiol Scand 117, 307–309.

Mancini, M., Salarian, A., Carlson-Kuhta, P., Zampieri, C., King, L., Chiari, L., and Horak, F.B. (2012). ISway: a sensitive, valid and reliable measure of postural control. J Neuroeng Rehabil 9, 59.

Mergner, T., and Rosemeier, T. (1998). Interaction of vestibular, somatosensory and visual signals for postural control and motion perception under terrestrial and microgravity conditions--a conceptual model. Brain Res Brain Res Rev 28, 118–135.

Moe-Nilssen, R. (1998). A new method for evaluating motor control in gait under real-life environmental conditions. Part 1: The instrument. Clin Biomech (Bristol, Avon) 13, 320–327.

Nicholas, S.C., Doxey-Gasway, D.D., and Paloski, W.H. (1998). A link-segment model of upright human posture for analysis of head-trunk coordination. J Vestib Res 8, 187–200.

Paloski, W.H., Wood, S.J., Feiveson, A.H., Black, F.O., Hwang, E.Y., and Reschke, M.F. (2006). Destabilization of human balance control by static and dynamic head tilts. Gait Posture 23, 315–323.

Pettorossi, V.E., and Schieppati, M. (2014). Neck proprioception shapes body orientation and perception of motion. Front Hum Neurosci 8, 895.

Pozzo, T., Berthoz, A., and Lefort, L. (1990). Head stabilization during various locomotor tasks in humans. I. Normal subjects. Exp Brain Res 82, 97–106.

Pozzo, T., Berthoz, A., Lefort, L., and Vitte, E. (1991). Head stabilization during various locomotor tasks in humans. II. Patients with bilateral peripheral vestibular deficits. Exp Brain Res 85, 208–217.

Pozzo, T., Levik, Y., and Berthoz, A. (1995). Head and trunk movements in the frontal plane during complex dynamic equilibrium tasks in humans. Exp Brain Res 106, 327–338.

Sakaguchi, M., Taguchi, K., Ishiyama, T., Netsu, K., and Sato, K. (1995). Relationship between head sway and center of foot pressure sway. Auris Nasus Larynx 22, 151–157.

Schmidt, J.D., Terry, D.P., Ko, J., Newell, K.M., and Miller, L.S. (2018). Balance Regularity Among Former High School Football Players With or Without a History of Concussion. J Athl Train 53, 109–114.

Scoppa, F., Capra, R., Gallamini, M., and Shiffer, R. (2013). Clinical stabilometry standardization: basic definitions--acquisition interval--sampling frequency. Gait Posture 37, 290–292.

Sessoms, P.H., Gottshall, K.R., Sturdy, J., and Viirre, E. (2015). Head Stabilization Measurements as a Potential Evaluation Tool for Comparison of Persons With TBI and Vestibular Dysfunction With Healthy Controls. Military Medicine 180, 135–142.

Shumway-Cook, A., and Horak, F.B. (1986). Assessing the influence of sensory interaction of balance. Suggestion from the field. Phys Ther 66, 1548–1550.

Sosnoff, J.J., Broglio, S.P., Shin, S., and Ferrara, M.S. (2011). Previous mild traumatic brain injury and postural-control dynamics. J Athl Train 46, 85–91.

Stoffregen, T.A., Hettinger, L.J., Haas, M.W., Roe, M.M., and Smart, L.J. (2000). Postural instability and motion sickness in a fixed-based flight simulator. Hum Factors 42, 458–469.

Woodson, J. (2015). “Traumatic Brain Injury: Updated Definition and Reporting”, (ed.) D.O. Defense. (Washington, DC).

Zatsiorsky, V.M., and Duarte, M. (1999). Instant equilibrium point and its migration in standing tasks: rambling and trembling components of the stabilogram. Motor control 3, 28–38.

Zhang, Y., Kiemel, T., and Jeka, J. (2007). The influence of sensory information on two-component coordination during quiet stance. Gait Posture 26, 263–271.

